# Development of Multi-Bundle Virtual Ligaments to Simulate Knee Mechanics after Total Knee Arthroplasty

**DOI:** 10.1101/2022.10.12.511986

**Authors:** Samira Vakili, Brent Lanting, Alan Getgood, Ryan Willing

## Abstract

Preclinical evaluation of total knee arthroplasty (TKA) components is essential to understanding their mechanical behavior and developing strategies for improving joint stability. While preclinical testing of TKA components has been useful in quantifying their effectiveness, such testing can be criticized for lacking clinical relevance, as the important contributions of surrounding soft tissues are either neglected or greatly simplified. The purpose of our study was to develop and determine if subject-specific virtual ligaments reproduce the same kinematics as native ligaments surrounding TKA joints. Five TKA knees were mounted to a motion simulator. Each was subjected to tests of anterior-posterior (AP), internal-external (IE), and varus-valgus (VV) laxity. The forces transmitted through major ligaments were measured using a sequential resection technique. By tuning the measured ligament forces and elongations to a generic non-linear elastic ligament model, virtual ligaments were designed and used to simulate the soft tissue envelope around isolated TKA components. The average root mean square error (RMSE) between the laxity results of TKA joints with native versus virtual ligaments was 2.9 mm during AP translation, 6.5° during IE rotations, and 2.0° during VV rotations, and there was no statistically significant difference between the results of both methods. Interclass correlation coefficients (ICCs) indicated a good level of reliability for AP and IE laxity (0.85 and 0.84). To conclude, a virtual ligament envelope around TKA joints can mimic natural knee behavior and is an effective method for the preclinical testing of TKA components.

## Introduction

Total knee arthroplasty (TKA) is a treatment for patients with end-stage knee osteoarthritis [1]. TKA aims to relieve pain, improve joint kinematics, and restore joint stability [2–7]. Instability is an often-cited reason for revision and patient dissatisfaction, and therefore an important aspect to consider when designing surgical procedures and TKA prosthesis components [8–12]. Preclinical testing of TKA components is essential to understand their mechanical performance and for developing strategies to improve joint stability [13]. Ideally, such preclinical tests should reproduce the physiological behavior of the knee joint, which is guided by the articular geometries of the joint in conjunction with the capsule and ligaments.

A new TKA design can be evaluated experimentally, computationally, and clinically [2,3,6,14–19]. While clinical outcomes are the gold standard, it can be hard to measure the effects of variability of a particular structure in isolation (e.g., ligament resection); moreover, designing prospective clinical studies of a new TKA system is costly and should only occur at the final stages of development. Thus, the focus of many experimental and computational methods is to replicate the *in vivo* environment that exists for a TKA components in the human body, as well as the roles that ligaments play in providing joint stability. Accurate replication of these surrounding mechanical structures is crucial for ensuring that the results of these studies are reliable. Computational studies are appealing for their reduced cost and utility for large-scale parametric analyses but incorporate a myriad of assumptions and require careful validation against experimental results [20–23]. The limitations and assumptions include the given simplified material properties and geometries, as well as the difficulty in creating and validating subject-specific models that match experimental results [24–28]. Furthermore, even in the case of simulating a subject-specific model with corresponding one-to-one experiments, many models do not consider the real ligament recruitment patterns present during knee motions, but instead only focus on replicating kinematics to achieve optimal joint stability [26,29]. It is possible these simplifications result in unrealistic forces and exhibit inconsistent and unrealistic results. Therefore, it is important to investigate methods to incorporate more realistic physiological loads and surrounding mechanical structures when assessing TKA joint stability.

Experiments utilizing cadaveric specimens on joint motion simulators offer the benefit of a realistic loading environment including surrounding tissues which contribute to knee joint motion and stability; however, these studies are expensive to extend beyond a small number of specimens and are not easily parameterized. Experiments using isolated prosthesis components on a joint motion simulator are useful for studying prosthesis behavior, but until now have lacked an accurate representation of the surrounding soft tissue envelop, which can influence measured joint kinematics and stability [30]. This shortcoming has recently been addressed by the development of joint motion simulators which can incorporate the mechanical contributions of virtual ligaments (virtual one-dimensional (1-D) point-to-point nonlinear elastic springs). The utility of these virtual ligaments has been demonstrated through previous studies of elbows and knees [18,31]. This enables the use of virtual ligaments as a replacement for native ligaments to enhance the accuracy of *in vitro* testing of isolated TKA prosthesis components on a joint motion simulator. While the ability to incorporate their behaviors has been provided, the challenge which remains is to properly define their mechanical properties to produce a virtual soft tissue envelope that behaves in a physiologically accurate manner.

Thus, the objective of this study was to characterize the soft tissue behaviors of native human knee ligaments *in situ* after a total knee arthroplasty, and then to re-apply these as virtual ligaments around isolated TKA prosthesis components on a joint motion simulator to validate their accuracy for the purpose of pre-clinical laxity testing of TKA prosthesis components.

## Method

### 1. Specimen preparation and TKA surgery

Five fresh frozen intact human cadaveric knee specimens (three males, three females, average age of 61 ± 10 and average BMI of 26 ± 6) were tested. Each frozen knee was CT scanned prior to the study and thawed for 24 hours at room temperature. Triathlon TKA components (Stryker Corp., Mahwah, NJ) were implanted into the knee joint by an orthopedic surgeon, in mechanical alignment using conventional surgical tools [32]. The joints were sized and balanced for cruciate-retaining ultra-high molecular weight polyethylene (UHMWPE) bearings. Each specimen was then mounted onto a 6 degrees-of-freedom (6-dof) joint motion simulator (AMTI VIVO, Advanced Mechanical Technology, Inc, Watertown, MA). The joint motion simulator provides displacement-control motion with a nominal positional resolution of 0.1 mm or 0.1° and force-control loading with less than 10 N or 0.5 Nm error (Figure 1A).

**Figure 1:**
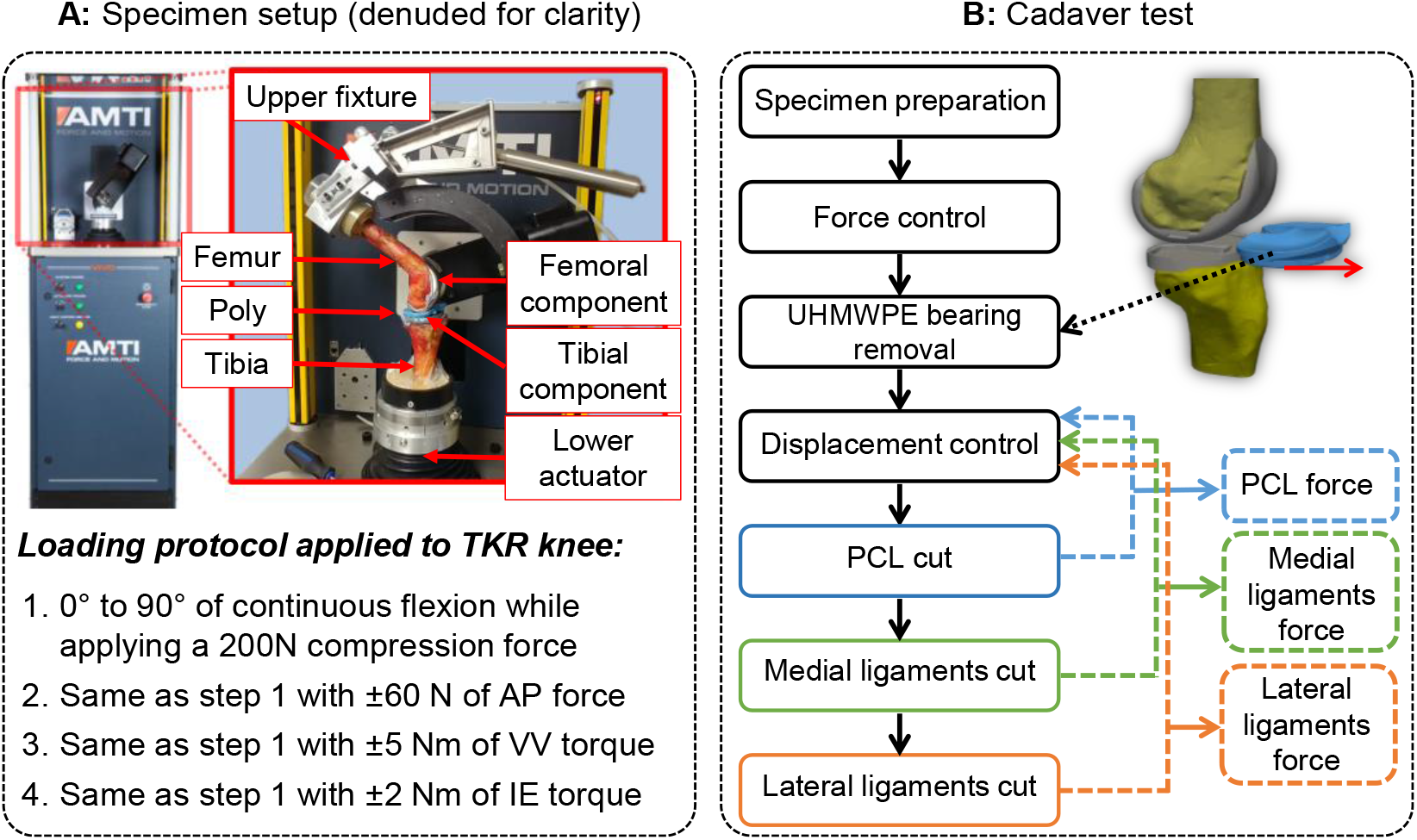
(A) Specimen setup on 6-dof AMTI VIVO joint motion simulator (joint denuded for clarity). (B) Cutting sequence for ligament force measurement

Using a custom-designed aluminum fixture, the proximal femur was potted and mounted onto the upper actuator of the simulator, such that the femoral flexion axis was aligned with the mechanical flexion axis of the apparatus. The tibia and fibula were then potted and secured to the lower actuator of the simulator. This potting technique minimized the amount of tibial secondary motion throughout knee flexion and has been described in greater detail in a previous study [6]. Then, a reference pose of the knee was determined by applying compression to the joint in full extension.

### 2. Simulated Motions

Each knee was subjected to loads that simulate knee laxity tests. Knee laxity tests were simulated by applying anterior-posterior (AP) forces, varus-valgus (VV), and internal-external (IE) torques (±60 N, ±5 Nm, or ±2 Nm, respectively) to the tibia while maintaining 200 N of compression, across the full range of flexion and extension between 0° and 90°.

### 3. Testing sequence

The loading scenarios described above were completed for each specimen and the resulting joint kinematics were measured. Subsequently, the UHMWPE bearing was removed, and the previously measured kinematics were re-applied to the knee in displacement control mode, and the resulting reaction forces and torques were recorded by a loadcell embedded in the tibial actuator. With the bearing removed, there was no contact between the femur and the tibia during the displacement-controlled joint motions, and all forces measured were those transmitted through soft tissues (Figure 1 B) [33]. To measure ligament tensions throughout each motion studied, we employed a sequential resection technique whereby each ligament was individually released, the joint was translated/rotated through the previously measured kinematics in displacement control, and changes in reaction forces were recorded. Using the principle of superposition, changes in the reaction forces and torques were directly related to the tension which was present in the ligament previously dissected [31,33,34]. This process was completed after resection of (1) the PCL, (2) all ligaments on the medial side of the knee, and (3) all ligaments on the lateral side of the knee (Figure 1B).

### 4. Ligament attachment sites prediction

After all the tests were completed, all remaining soft tissues were denuded, and the bone (femur and tibia) and TKA prosthesis component surfaces were digitized using an articulating arm digital coordinate measuring machine (GAGE, FARO Technologies, Lake Mary, FL, USA) with the specimen maintained in the pre-determined reference pose on the motion simulator. Three-dimensional (3-D) reconstructions of the femur, tibia and fibula were created in 3-D Slicer (http://www.slicer.org/) using the intact knee CT scans images. Using a surface-based registration technique [35], these 3-D surface models were co-registered to their corresponding surface digitization. This allowed for characterization of prosthesis component alignments, positions of ligament insertions with respect to TKA components and bone to measure ligament elongation during motions.

### 5. Optimization

An optimization technique was used to characterize the biomechanical properties of five major knee ligaments crossing the tibiofemoral joint, including the posterior cruciate ligament (PCL), the lateral collateral ligament (LCL), the anterolateral ligament (ALL), the posterior oblique ligament (POL) and the superficial bundle of the medial collateral ligament (sMCL). The anterior cruciate ligament (ACL) was not considered since it was dissected during TKA. The ligament insertion and origin points of the PCL bundle groups (the anterolateral PCL (alPCL) and posteromedial PCL (pmPCL)), LCL, ALL, POL and sMCL were determined in one of the knee models (as a template model) based on anatomical descriptions in the literature [35– 37] (Figure 2). The reference model with embedded ligament attachment regions was then morphed to other knee models using a non-rigid iterative closest point method in MATLAB (MathWorks, Natick, MA) [38–42]. This allowed us to map the template insertions onto each specimen for subject-specific ligament attachment regions.

**Figure 2:**
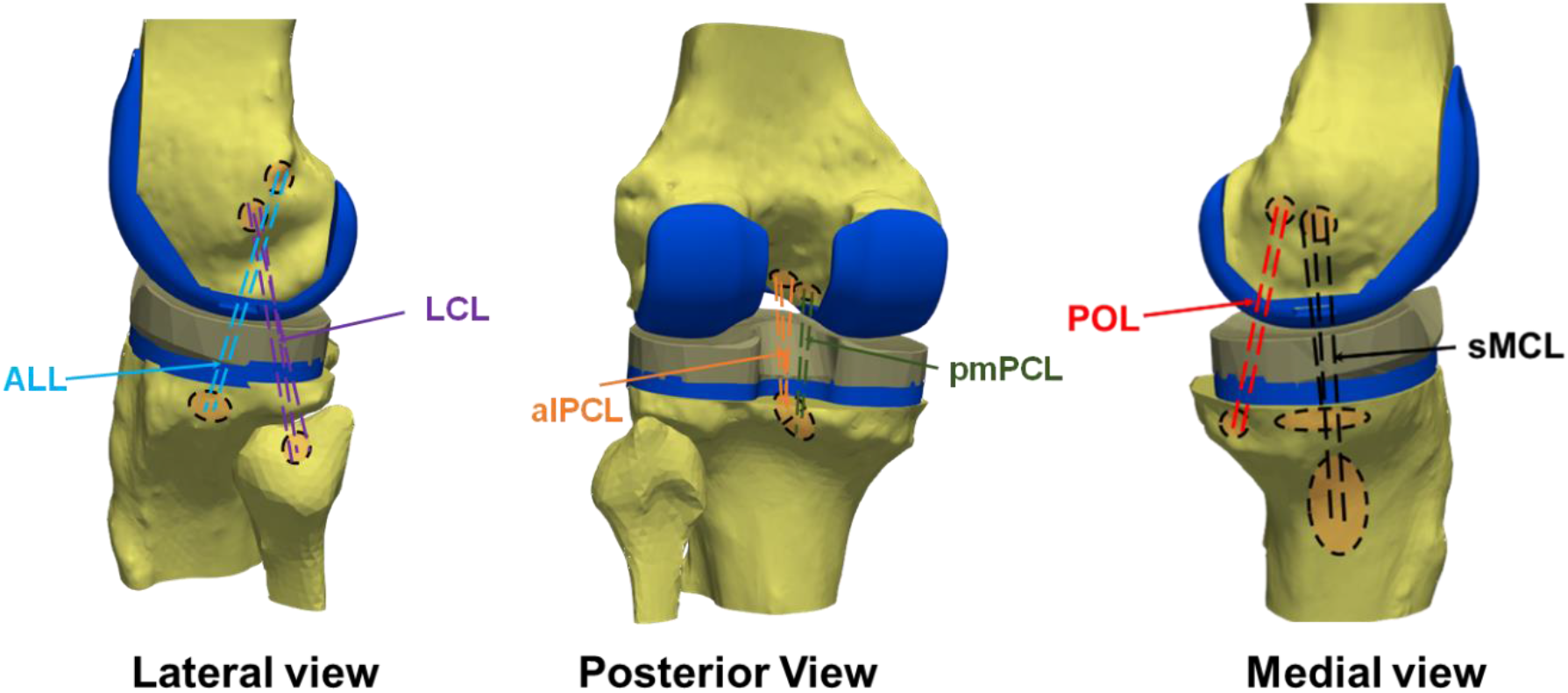
Lateral, posterior and medial view of a 3-D model of the left knee joint constructed from the CT scan images. The lines connecting the attachment sites represent the posterior cruciate ligament (two bundle groups- the anterolateral PCL (alPCL) and posteromedial PCL (pmPCL)), the lateral collateral ligament (LCL), the anterolateral ligament (ALL), the posterior oblique ligament (POL) and the superficial bundle of the medial collateral ligament (sMCL).

Ligament lines of action and elongations were computed based on the relative positions of the femoral and tibial insertions of each bundle in the determined ligament’s attachment area, according to the measured kinematics for each knee during all motions. The patient-specific mechanical properties of each ligament were characterized using an established nonlinear force-strain relationship (Eq. 1) [43–45], where k is stiffness and ε is ligament strain, l_slack_ is the slack length of the ligament at which the ligament begins to transmit tensile force, and ε_l_ is the spring parameter assumed to be 0.03 [46].

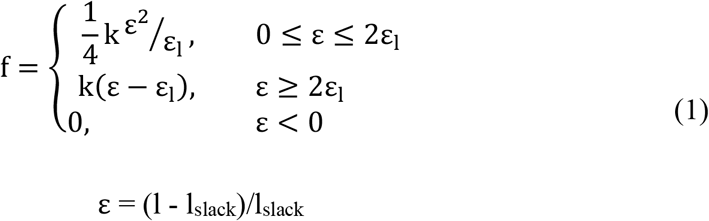

Using multi-objective optimization in MATLAB’s global optimization toolbox, ligament mechanical properties, including k, reference strain (ε_0_), the number of bundles per ligament, and their insertion coordinates were optimized. ε_0_ was associated with the measured initial ligament strain in the pre-determined reference pose which is defined by ε_0_ = (l_0_ - l_slack_)/l_slack_, where l_0_ is the ligament length in the pre-determined reference pose. Multi-objective optimization was conducted using three cost functions for minimizing the root mean square errors (RMSE) between calculated and experimentally measured ligament forces in the anterior-posterior (AP), medial-lateral (ML), and superior-inferior (SI) directions. During the optimization process, ligament attachment locations were allowed to vary within the distribution of the ligament’s predicted attachment area. The stiffness parameter was allowed to vary up to 50% lower than the value that was reported in the literature and no upper limit was imposed [46,47]. The stiffness parameter of each ligament group was set so that all bundles within a ligament division had the same stiffness but different slack lengths. To maintain ligament strains within a physiological range, the maximum allowable ligament strain across all motions during the optimization process was constrained to be less than 10% [48,49]. The number of ligament bundles of the alPCL, pmPCL, LCL, ALL, POL and sMCL varied to a maximum of 5 bundles for each ligament group. The final set of virtual ligaments was determined among the Pareto front results based on the minimum RMSE between experimental and calculated force in AP, ML and SI direction. A brief overview of the design optimization process is shown in Figure 3.

**Figure 3:**
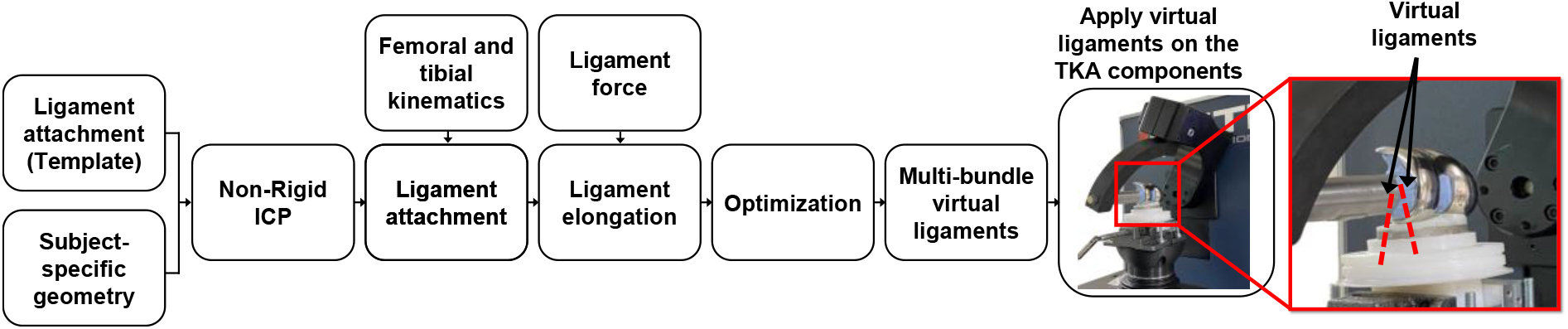
The framework for designing virtual ligaments, and implementing the designed virtual ligaments on TKA prosthesis components to measure joint laxity limits

### 6. Virtual ligaments

Virtual ligaments connect the femoral and tibial attachment points with tension-only non-linear virtual springs, which can replicate the continuous nonlinear force-elongation response of native ligaments entirely within the joint motion simulator’s control system [18,31]. We performed hybrid experimental-computational testing on the joint motion simulator using the same TKA prosthesis components that were used during cadaveric testing, with each of the optimized virtual ligament sets, by providing the femoral and tibial attachment sites and material properties (k, ε_0_) for each bundle as measured before (Figure 3). Virtual ligaments were applied to the knee prosthesis components in their same relative positions to tibia and femur as they were in the experiment. The same loads were reapplied, which allowed the comparison of joint behaviors resulting from a TKA stabilized by native ligaments in a cadaver knee versus the same TKA mounted directly to the motion simulator and stabilized by virtual ligaments. The laxity kinematics were measured in the AP, IE and VV directions at 0°, 30°, 60°, and 90° of flexion.

### 7. Data analysis

For both cadaver and isolated TKA prosthesis component tests with virtual ligaments, kinematic data was filtered with a low-pass Butterworth filter and then down-sampled to 1024 samples per cycle using the spline interpolation function in MATLAB. The kinematics of the knee were then sampled in 15° increments of flexion from 0° to 90°, and the average of the flexion/extension data at each angle was calculated. The output variables were the AP motion limits during anterior and posterior directed force loading, the IE motion limits during internal and external torque loading, and the VV motion limits during varus and valgus torque loading. AP, IE and VV laxities were calculated as the differences between positive and negative AP, IE and VV motion limits.

Statistical analyses were performed using a repeated measures analysis of variance (ANOVA) with a Greenhouse-Geisser correction in SPSS (IBM Corp., Armonk, NY, USA) to determine if the method of testing of TKA prosthetic components (stabilized with native versus virtual ligaments) in different flexion angles (0°–90° in 15° increments) had significant effects on AP, IE, and VV laxity limits. The effects of method of testing on AP, IE, and VV laxity were compared using post hoc comparisons of means with a Bonferroni correction with a threshold of 0.05 where significant differences were detected.

The agreement between the results of the joints with native and virtual ligaments was further investigated using a Bland-Altman analysis [50]. The method’s accuracy is determined by its bias between the mean differences, while its precision was determined by its limit of agreement. The limit of agreement was set at 1.96 standard deviations (SDs), providing an interval within which 95% of differences between kinematics of joints with native ligaments and those with virtual ligaments were expected to occur. Finally, the Interclass correlation coefficient (ICC) was conducted to measure the reliability of testing TKA joints with virtual ligaments compared to native ligaments, using a two-way random effects model for absolute agreement in SPSS. An ICC value ranges from 0 to 1, with values below 0.5 indicating poor reliability, between 0.5 and 0.75 moderate reliability, between 0.75 and 0.9 good reliability, and above 0.9 excellent reliability [51].

## Results

Figure 4 shows the AP laxity of five TKA joints with native ligaments and virtual ligaments, along with their mean and SD. The RMSE between the AP laxity for the joint with native versus virtual ligaments was 2.1 ± 1.1 mm, 2.9 ± 1.7 mm, 3.1 ± 2.0 mm, and 3.7 ± 1.5 mm at 0°, 30°, 60°, and 90° of flexion, respectively. AP laxity results did not show any significant difference when using virtual versus native ligaments (p = 0.3), as well as no effect on the results of using different method of testing in different flexion angles (p = 0.2).

**Figure 4:**
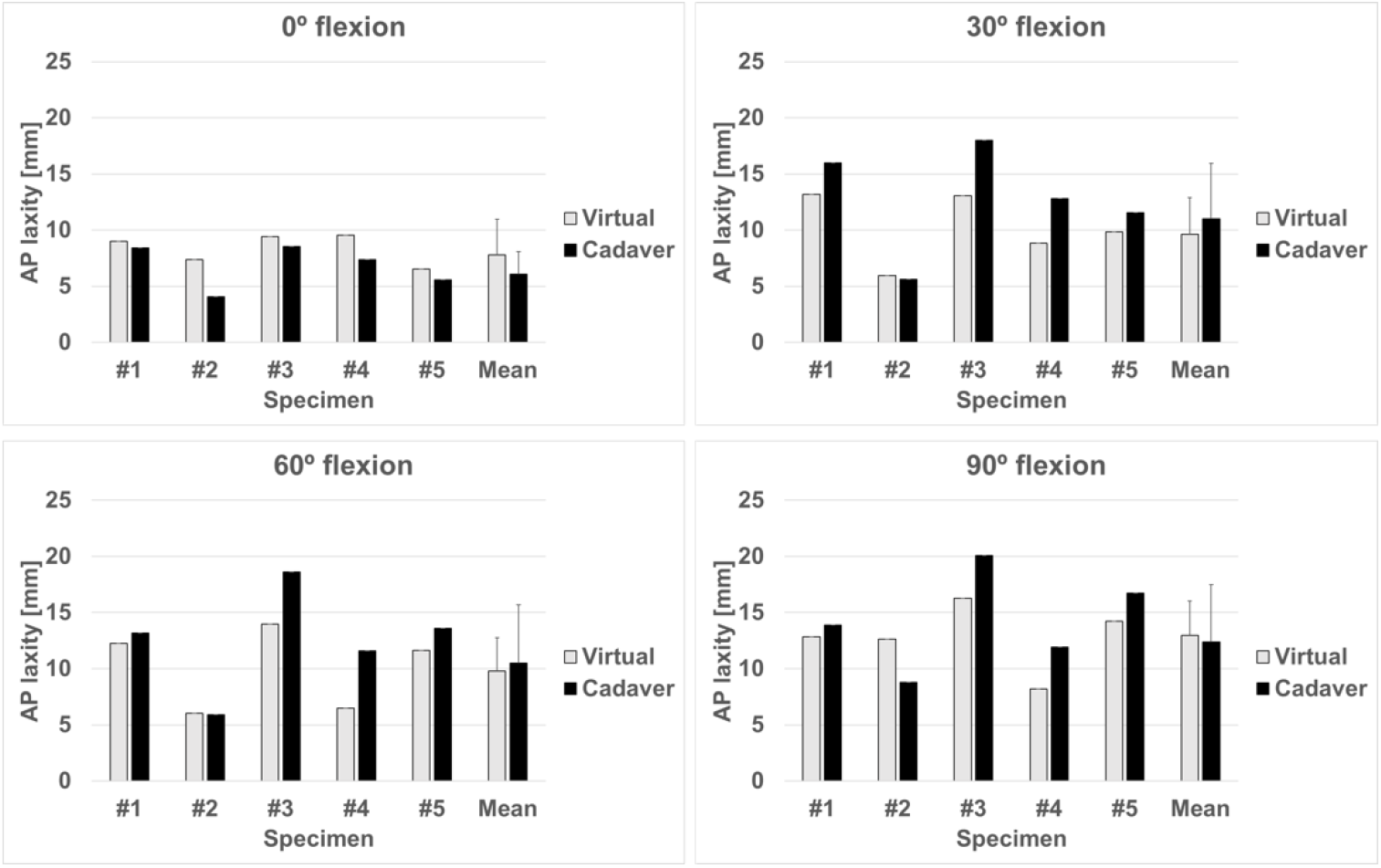
AP laxity in TKA joint with native ligaments and with virtual ligaments tests in 0°, 30°, 60° and 90° of flexion for each specimen, as well as the mean ± SD.

The IE laxity for five TKA joints with native and virtual ligaments, along with their mean and SD. are shown in Figure 5. The RMSE between the IE laxity for the joint with native versus virtual ligaments was also 4.6 ± 1.7°, 6.3 ± 3.6°, 7.0 ± 3.2°, and 7.7 ± 5.1°, for 0°, 30°, 60°, and 90° of flexion angle. IE laxity results did not differ statistically significant when using virtual versus native ligaments (p = 0.2), as well as no interaction between the method of testing in different flexion angles (p = 0.7).

**Figure 5:**
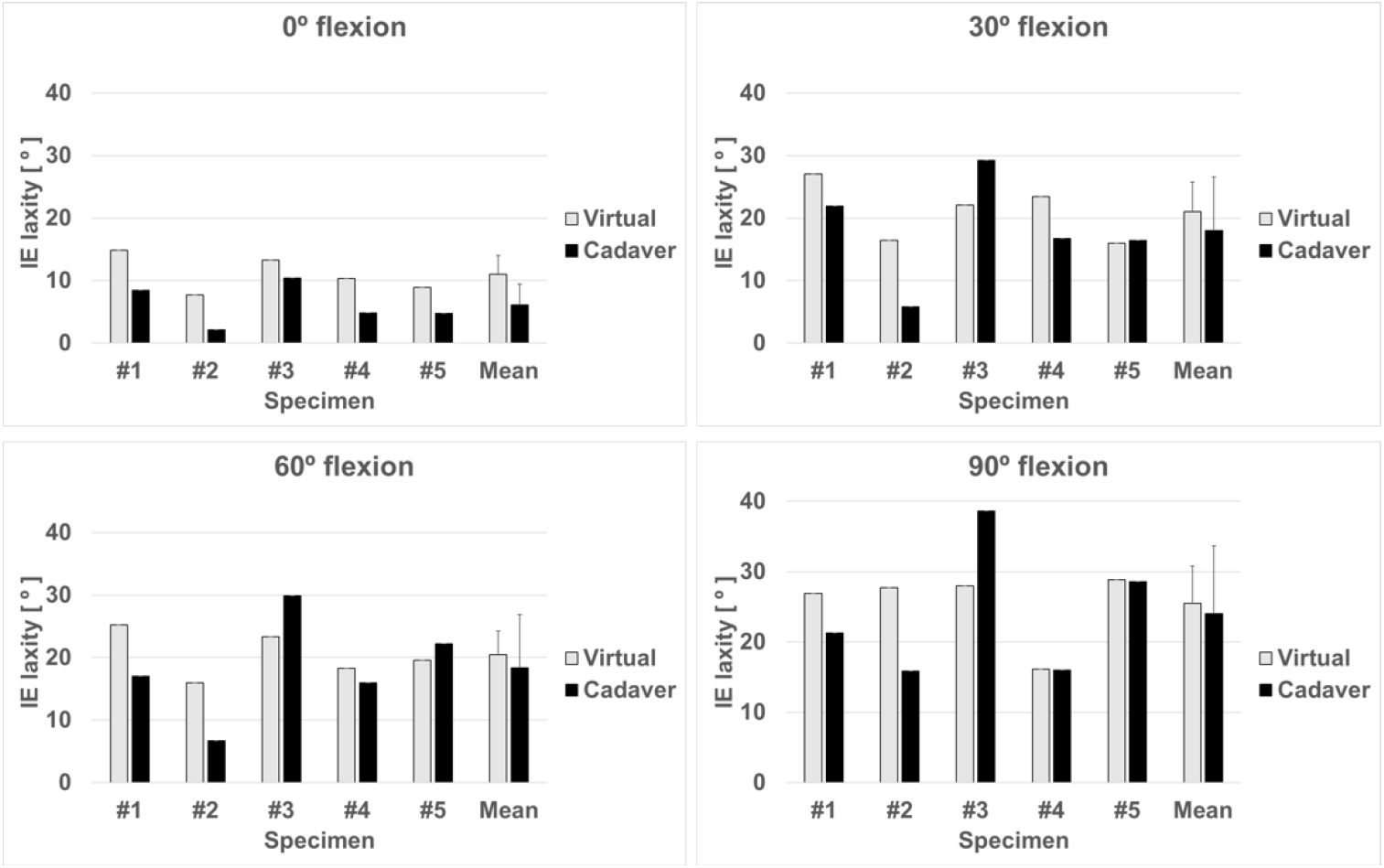
IE laxity in TKA joint with native ligaments and with virtual ligaments tests in 0°, 30°, 60° and 90° of flexion for each specimen, as well as the mean ± SD.

The VV laxity for five TKA joints with native and virtual ligaments, along with their mean and SD are shown in Figure 6. The RMSE between the VV laxity for the joint with native versus virtual ligaments was 0.7 ± 0.4°, 2.7 ± 1.5°, 2.4 ± 1.7°, and 2.5 ± 1.2°, for 0°, 30°, 60°, and 90° of flexion angle. The results of VV laxity for the joints with virtual ligaments were not statistically significant different when compared to the joints with native ligaments (p = 0.1), as well as no interaction between the method of testing in different flexion angles (p = 0.4).

**Figure 6:**
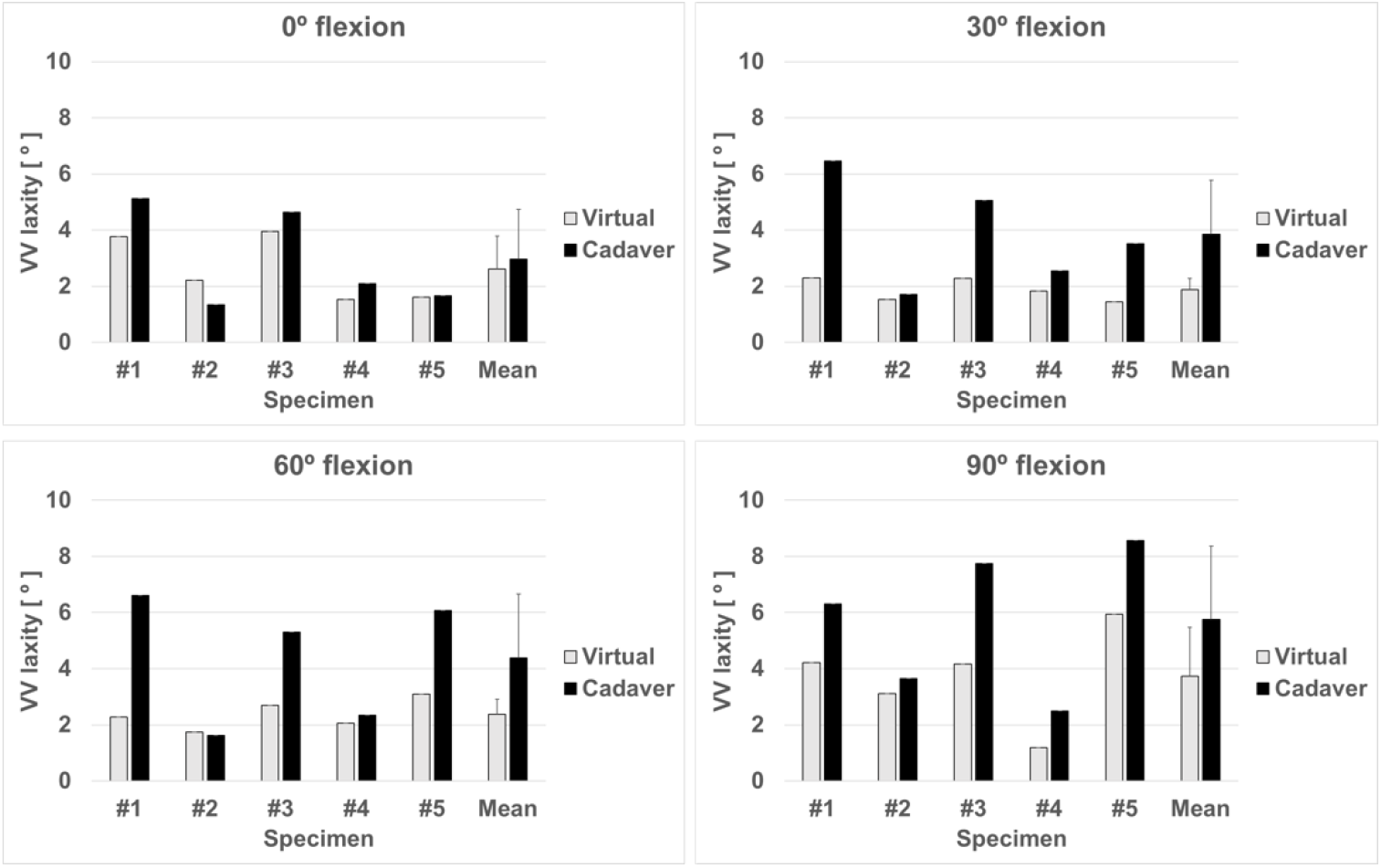
VV laxity in TKA joint with native ligaments and with virtual ligaments tests in 0°, 30°, 60° and 90° of flexion for each specimen, as well as the mean ± SD.

The agreement between the results of the joints with native and virtual ligaments was further investigated using a Bland-Altman analysis [50] as shown in Figure 7 A. The method’s accuracy is determined by its bias, while its precision was determined by its limit of agreement. Bland-Altman plots showed a bias of 0.9 mm, -3.1° and 1.1° for AP, IE and VV laxity between the results of two methods, respectively. The limit of agreement was 6.6 mm and -4.0 mm in the AP laxity, 8.9° and -14.6° in the IE laxity, and 4.5° and -1.4° in the VV laxity. In Figure 7 B, the scatter plot of laxity limits of TKA joints with native versus virtual ligaments is plotted along with a line of equality in order to demonstrate the correlation between both techniques. These plots indicate that AP and IE laxities using virtual ligaments are overpredicted at smaller laxities, while they are underpredicted at higher laxities. As a result of using virtual ligaments, VV laxities are generally underestimated when compared with cadaver tests.

**Figure 7:**
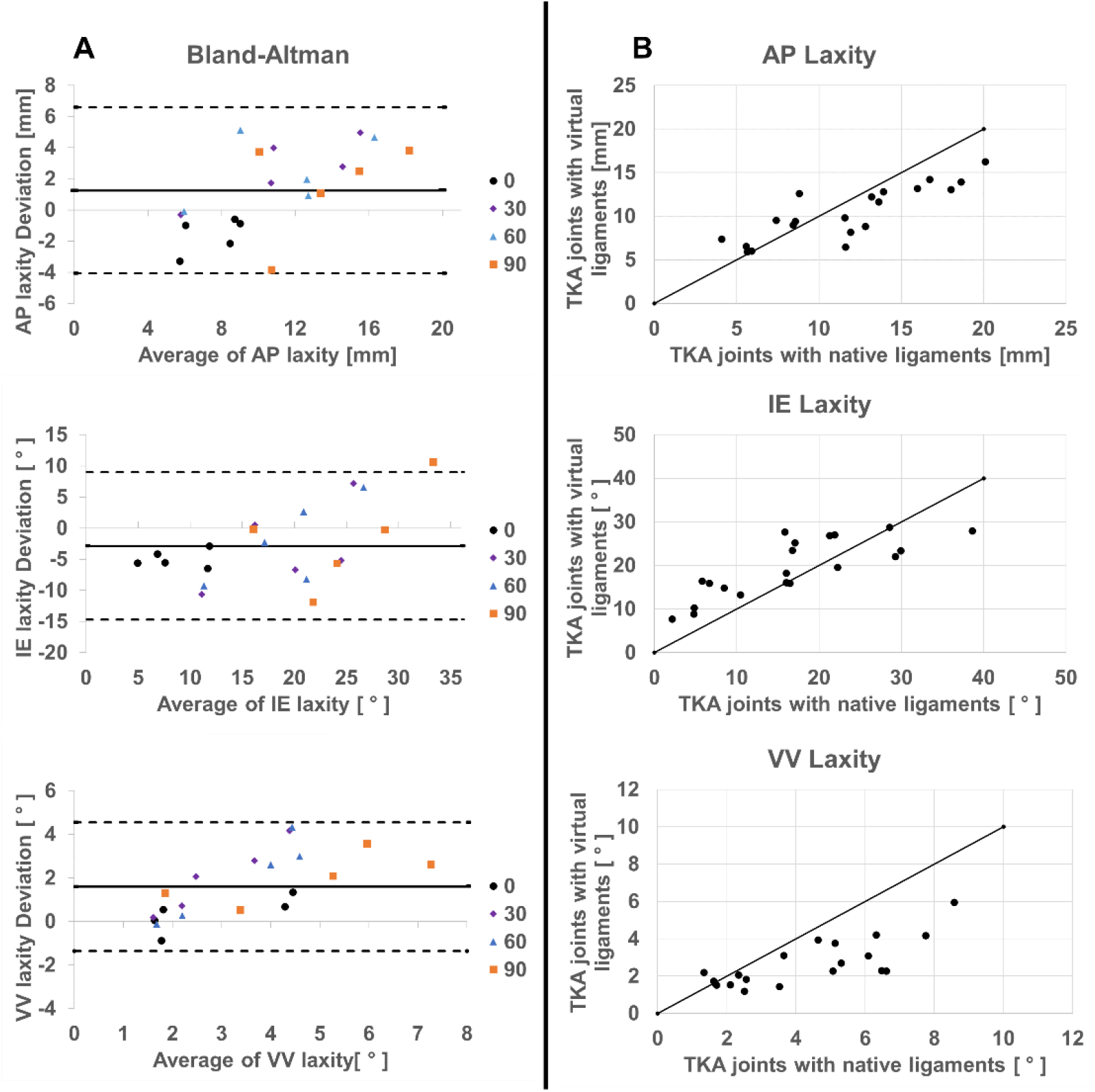
(A) Bland–Altman plots comparing the measurements from the joints with native ligaments and with those of virtual ligaments for measuring AP, IE and VV laxities. In this plots, accurate laxity results showed a bias (continuous line) close to the line ‘x=0’, whereas precision showed limits of agreement close to the bias (dashed lines). (B) Scatter plots for laxity measurements of TKA joints with native ligaments on the horizontal axis and with virtual ligaments on the vertical axis. The line of equality is shown with a black solid line.

ICC also was conducted to measure the reliability of testing TKA joints with virtual ligament compared to native ligaments [51]. ICC was 0.85 and 0.84 for AP and IE laxity, respectively, indicated a good level of reliability between both methods, while VV laxity results showed moderate reliability (ICC = 0.66).

## Discussion

The purpose of this study was to develop a combined experimental and computational framework for characterizing the force and elongation contributions of knee native ligaments to simulate virtual ligaments that could be used as a substitute for native ligaments during pre-clinical evaluations of TKA joint stability. We calibrated the virtual ligament load-elongation behavior by using a generic ligament model to represent the ligaments’ recruitment during knee motions that leads to stabilizing TKA joints in laxity motions. Our virtual ligament design approach, including calibration of ligament parameters, was similar to the methods of previous studies [31,45]. However, in the present study, we have provided a more constrained multi-objective optimization than prior studies by directly calibrating the subject-specific virtual ligaments properties for five cadaveric TKA joints throughout a continuous flexion range of motion while applying superimposed force and torques during laxity tests.

The average RMSE values between TKA joints with native versus virtual ligaments for the laxity tests in the current study was 2.9 mm during AP translation, 6.5° during IE rotations, and 2.0° during VV rotations. The RMSE between the kinematics of the joints during native versus virtual ligament tests was similar to Harris et al.’s findings for a finite element model that minimized both kinematics and ligament recruitment during knee motions between their models and experiment results [29]. Harris et al., used a combined experimental and computational technique to analyze the laxity of knee joints by providing subject-specific experimental kinematics for calibrating a finite element model whilst incorporating ligament recruitment during knee motions into the model calibration based on literature. In their study, errors between model-predicted and experimental kinematics averaged less than 3 mm of translation during AP displacements, less than 6° during IE rotations, and less than 2° during VV rotations. Their findings emphasize the importance of considering ligament engagement when calibrating knee models, which was the focus of our study. The present study also showed similar errors to those reported by Blankevoort et al., when their 3-D mathematical models of the tibiofemoral joint were loaded with 3 Nm of internal torque and 100 N of AP load [26]. They report 0 to 8° of error in internal rotations and 3 to 5 mm of error in AP displacements. Baldwin et al., also report the average 2.4° and 0.6° of IE and VV errors reported for TKA knees subjected to similar loading profiles [20]. Both Blankevoort et al. and Baldwin et al. studies aimed to minimize the difference between their mathematical model and the experiment’s kinematics without taking ligament recruitment into account. There were similar errors in our study as in the other studies, but the errors in our study were slightly greater than those observed in finite element modeling studies [20,26,29]. It is important to note that the optimization in computational modeling studies differs from ours, as their model ligament properties were calibrated to minimize the difference between the model and experimental kinematics, resulting in a smaller kinematic error. In contrast, the optimization in our study aimed to tune the ligament properties to simulate the load-elongation behavior of the ligaments surrounding the knee, and then applying those ligaments to the joint in order to determine the effectiveness of the optimization based on the resulting kinematics of the joint. This is a more challenging indirect optimization approach.

The Bland-Altman analysis was also conducted to compare the mean differences and the limits of agreement between the result of TKA joint with native versus virtual ligaments, as well as the scatter plot with the line of equality to demonstrate a clear trend between the two methods (Figure 7). These results indicate that there is a fixed bias between the laxity results of the two methods in all laxity limits. A smaller bias was observed for AP laxity while a larger bias was observed for IE laxity. For AP, IE, and VV laxities, the limit of agreement was within an acceptable range. Although there was no direct comparison in the literature to determine an acceptable limit of agreement, the average mean difference was comparable with the results of the computational study [20,26,29]. Furthermore, the results indicated that virtual ligaments overpredicted what should be small laxities and underpredicted what should be greater laxities, mainly in the AP and IE directions. This may be attributed to the fact that the ligament reference strain in our optimization was limited to be within the physiological range (10%) [49], thus the reference strain was smaller, the slack length was longer, and the stiffness was boosted to generate the ligament force, resulting in the ligaments engaging later but becoming stiffer following engagement.

Ultimately, to measure the reliability of testing TKA joints with virtual ligaments compared to native ligaments ICC was calculated. ICC indicated a good level of reliability between both methods for IE and AP laxities, while VV laxity results showed moderate reliability. The low level of agreement for VV laxity may be due to the fact that ligaments were modeled as 1D point-to-point spring elements and wrapping was not taken into account specially for collateral ligaments which are the main contributors in VV laxity [52].

There were limitations to both the TKA joints test with native ligaments (cadavers) and with virtual ligaments in the current study. During the cadaveric testing protocol, extra care was taken to maintain structural integrity of the knee during TKA and resection states; however, dissection of ligaments may have disrupted tissues surrounding the knee, such as the capsule. Consequently, the mechanical environment of the knee may had changed between each resection state, thus confounding ligament force results. Also, dissection of the entirety of ligaments on the medial and lateral sides makes it difficult to characterize individual ligament bundle recruitment on each side. Testing of TKA joints with virtual ligaments were also limited by the fact that simple 1-D springs were used to represent complex 3-D ligaments; however, the spring representations were computationally efficient in establishing the general constraint and force-elongation behavior of the ligaments [28]. Furthermore, the virtual ligaments were optimized and modelled for the entire range of flexion from 0° to 90°. There have been previous studies showing that optimizing ligament behavior at deeper flexion angles requires a separate set of ligament parameters [27] which explain the higher RMSE in higher flexion angles, most notably the IE rotation.

To the best of our knowledge, this was the first study to employ specimen-specific soft-tissue structures (virtual ligaments) to investigate the stability of TKA joints, which was the unique strength of this study. With virtual ligaments, the performance of TKA prosthesis components can be evaluated in future studies by comparing designs that preserve or sacrifice ligaments, as well as their performance during activities of daily living such as gait and stair climbing. Additionally, future work could focus on building a larger database of virtual knee ligaments, which could be used for preclinical testing of new devices across a “population” of virtual knee ligaments. Moreover, with virtual ligaments soft tissue models could be altered to simulate different pathologies or surgical approaches.

## Notes

### Competing Interest Statement

The authors have declared no competing interest.

